# Efficient septum formation is essential for chromosome segregation in *Bacillus subtilis* when SMC function is impaired

**DOI:** 10.64898/2026.06.22.733809

**Authors:** Ngoc Khanh Lai, Laura C. Lastra, Kehinde O. Adebiyi, David Z. Rudner, Stephen C. Jacobson, Daniel B. Kearns, Xindan Wang

## Abstract

Structural maintenance of chromosomes (SMC) complexes play conserved roles in chromosome organization, segregation, and repair in all domains of life. In *Bacillus subtilis*, SMC is required for segregation of newly replicated origins. To investigate whether other proteins function with SMC in this process, we performed a synthetic lethal screen with an *smc* hypomorphic allele (*smc*\*) that is mildly defective in chromosome segregation. In addition to recovering previously reported interactions of *smc* with *parB* and *spoIIIE*, our screen identified *minJ* and *divIVA* as essential in the *smc\** background. We show that the synthetic lethality between *smc\** and *ΔminJ* or *ΔdivIVA* arises from defects in segregating the replication terminus. Importantly, deletion of *minD,* which suppresses the cell division defects of *ΔminJ* and *ΔdivIVA*, restored terminus segregation and viability in the *smc\** background. These findings support a model in which proper septum formation promotes chromosome terminus resolution and segregation by enabling SpoIIIE-mediated DNA clearance from the division septum during cytokinesis. These findings highlight the interdependence between chromosome segregation and cell division.

**Importance:** The SMC complex plays a central role in chromosome organization and segregation in *Bacillus subtilis*, but the cellular functions that become important when SMC activity is reduced are not well understood. Using a hypomorphic *smc* allele, we discovered that mutations affecting cell division become essential when chromosome organization and segregation are impaired. Our findings support a model in which efficient septum formation enables proper localization of the SpoIIIE DNA translocase, which in turn resolves and segregates the chromosome terminus region. These results highlight the critical role of cell division in supporting chromosome segregation.

## Introduction

Bacterial chromosomes are compacted more than 1000-fold to fit within their cellular compartment (1, 2). Despite this compaction, the genome must be faithfully replicated and accurately segregated during each cell cycle. Maintaining the fidelity of these processes requires tight coordination between chromosome replication, segregation, and cell division.

In bacteria, the chromosome is segregated concurrently with DNA replication. Chromosome segregation is primarily mediated by the ParABS partitioning system and the structural maintenance of chromosomes (SMC) complexes (1, 2). The ParABS system is present in most bacterial genomes and plays a central role in chromosome segregation in many species (3, 4). ParB binds to *parS* sequences near the replication origin (4). Following replication initiation, ParB interacts with the ATPase ParA to drive bidirectional origin segregation (3, 5–7). ParB/*parS* complexes also serve as loading sites for SMC complexes (8–10), which are highly conserved chromosome organizers found across all domains of life (11–13). Once loaded near the origin, SMC complexes translocate towards the terminus, progressively aligning the left and right chromosome arms (14–19). Continuous loading and translocation of SMC complexes also promote sister chromosome resolution (20–22). The relative contributions of ParABS and SMC to chromosome segregation vary across species. For example, *Caulobacter crescentus* requires the ParABS system, whereas its SMC complex is dispensable (23, 24). In contrast, *Escherichia coli* lacks a ParABS system and instead relies on the SMC-related MukBEF complex for chromosome organization and segregation (25). Similarly, rapidly growing *B. subtilis* cells depend on SMC for viability, whereas the ParABS system is dispensable (26). Although *smc* mutants are viable at low growth rates, they exhibit defects in origin resolution and bulk chromosome segregation, resulting in elevated frequencies of anucleate cells (20, 21, 27–30).

At the final stage of segregation, the conserved FtsK/SpoIIIE family of DNA translocases resolves chromosome linkages at the divisome (31, 32). In *E. coli*, FtsK is recruited to the divisome during each cell division. Its C-terminal domain recognizes polarized chromosomal KOPS (FtsK-orienting polar sequences) motifs and directs DNA translocation toward the *dif* site, promoting chromosome segregation, dimer resolution, and decatenation (33–37). This activity clears DNA from the closing division plane prior to cytokinesis. In *B. subtilis*, the homologous SpoIIIE translocase performs a similar DNA clearance function. Most notably, during sporulation, SpoIIIE pumps chromosomal DNA across the asymmetric septum into the forespore, whereas during vegetative growth, it is largely dispensable and is recruited when the chromosomes become trapped at the division septum (38–41).

Chromosome segregation is closely linked to division-site selection in rod-shaped bacteria. Assembly of the division machinery begins with the formation of the FtsZ ring (Z-ring) at midcell (42–44). Proper Z-ring positioning is regulated by the Min system and nucleoid occlusion. In *B. subtilis*, the Min system is composed of four proteins, MinC, MinD, MinJ, and DivIVA, and functions both to prevent Z-ring formation near the cell poles and to promote Z-ring disassembly after septation (45–52). DivIVA localizes to regions of negative membrane curvature at the cell poles (53), where it recruits the transmembrane protein MinJ. MinJ interacts with the membrane-bound ATPase MinD, which then recruits the cell division inhibitor MinC to the membrane (42, 44, 49). MinC destabilizes FtsZ polymers, thereby inhibiting polar cell division (45, 49, 54). In parallel, nucleoid occlusion prevents Z-ring migration and assembly over the chromosome through the action of a DNA-binding protein that recognizes sequence elements largely absent from the replication terminus region (55–59). As chromosomes segregate toward the cell poles, the terminus region becomes positioned at midcell, relieving nucleoid occlusion and permitting Z-ring assembly and septum formation. Thus, divisome assembly and cytokinesis are tightly linked to chromosome segregation.

Genetic studies have revealed functional interactions between SMC and other chromosome segregation factors. Using a candidate-based approach, previous studies showed that cells lacking *smc* could not be transformed with either *ΔparB* or *ΔspoIIIE* (20, 60). Similarly, SMC-family proteins exhibit synthetic lethal interaction with FtsK in both *E. coli* and *Corynebacterium glutamicum* (61–63). These findings suggest that cells with impaired chromosome segregation become critically dependent on pathways that promote origin segregation and terminus resolution. To identify a more complete set of factors that become essential when chromosome segregation is impaired, we used transposon-sequencing (Tn-seq) to systematically screen for synthetic lethal partners of a hypomorphic *smc* allele (*smc\**). Consistent with previous studies (20, 60), we identified Δ*parB* and Δ*spoIIIE* as synthetic lethal with *smc*.* Our screen also identified Δ*minJ* and Δ*divIVA.* Cytological characterization of these mutants indicated that impaired chromosome segregation in *smc\** cells causes replicated termini to remain trapped at midcell. Under these conditions, SpoIIIE becomes essential for their resolution. Furthermore, defects in divisome assembly caused by the absence of *minJ* or *divIVA* prevent terminus resolution and result in a loss of viability.

## Results

### Isolation of a hypomorphic *smc* mutant (*smc\**)

To identify synthetic lethal partners of SMC, we sought to perform a genetic screen using an *smc* mutant. Because the *Δsmc* mutant is only viable at low temperature or on minimal medium and is prone to accumulating suppressors (27, 28), we isolated hypomorphic *smc* alleles for subsequent analysis. Using error-prone PCR, we generated a library of *smc* mutants at the endogenous *smc* locus (see Materials and Methods) and screened for mutants that form smaller colonies than wild-type (WT), indicative of partial-loss-of-function alleles of *smc*. Because *ΔparB* is a known synthetic lethal partner of SMC mutants (20, 21, 60), we screened the *smc* hypomorphs showing a loss of viability when transformed into *ΔparB*. From a set of candidates, we chose the isolate with the least severe growth defect to avoid bias toward general slow-growth phenotypes. This mutant contained a single amino acid substitution, V1119L, in SMC’s ATPase Walker B motif (11). For simplicity, we refer to *smc*(V1119L) as *smc\** (**Fig. 1A**).

**Figure 1.**
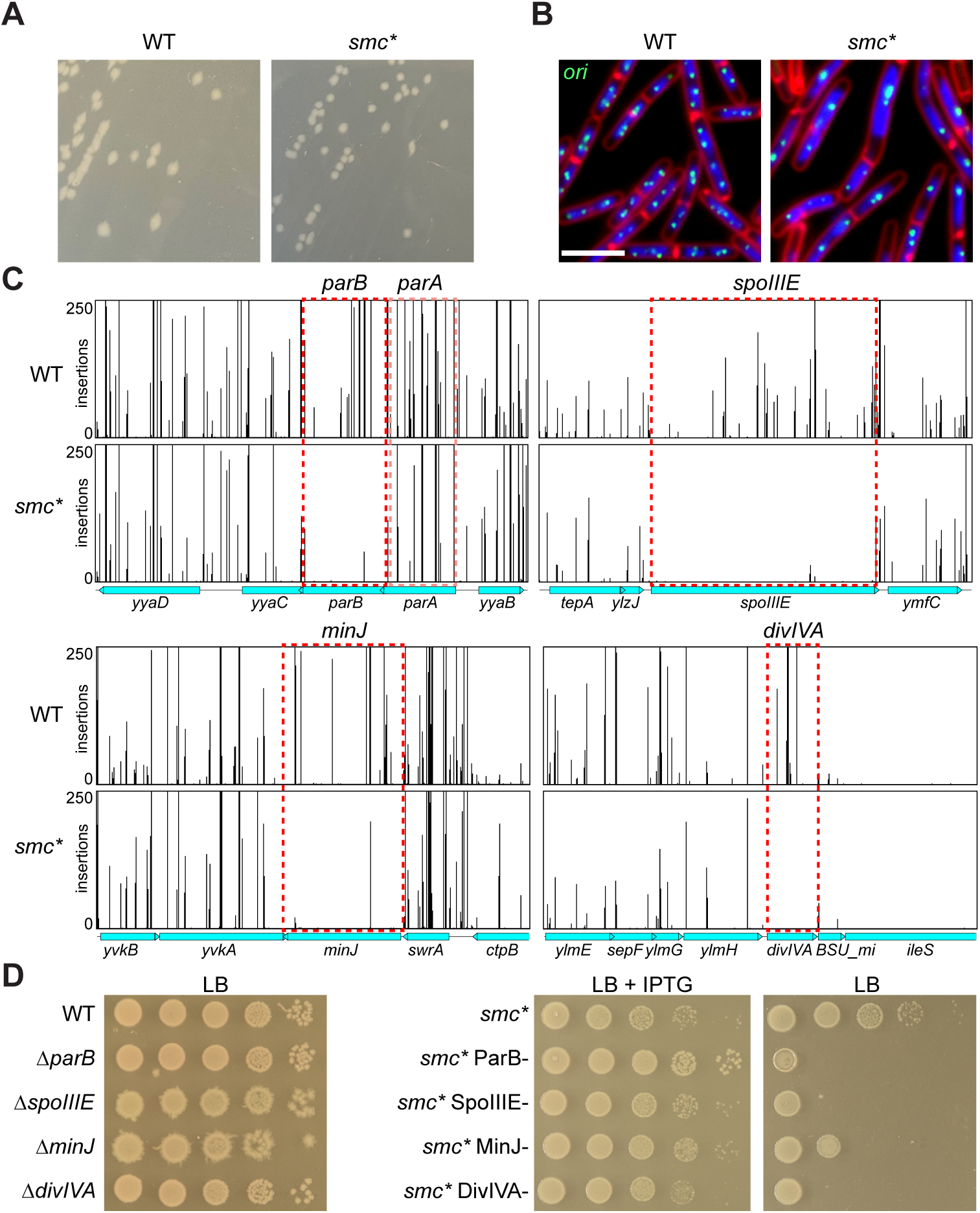
Isolation of a hypomorphic *smc\** allele and identification of synthetic lethal interactions. (A) WT (BWX5786, left) and *smc\** (BWX5790, right) colonies grown on an LB agar plate at 37°C. (B) Representative images of WT and *smc\** cells stained with FM4-64 to visualize membranes (red) and DAPI to visualize nucleoids (blue). Origins of replication were visualized using the *tetO*/TetR-CFP system (green). Scale bar, 4 μm. (C) Tn-seq plots showing transposon insertions at *parB*, *spoIIIE*, *minJ,* and *divIVA* in WT and *smc\** backgrounds. The x-axis indicates genomic position, and the y-axis indicates the number of transposon insertions at each insertion site. Dotted boxes highlight regions of indicated genes. (D) Serial dilutions of the indicated strains on LB agar plates with and without 0.5 mM IPTG. Left, as positive controls, WT (BWX5786), *ΔparB* (BWX5839), *ΔspoIIIE* (BWX5831), *ΔminJ* (BWX5837), *ΔdivIVA* (BWX5835) single-mutation cells were shown. Right, *smc\** (BWX5790), *smc\** ParB- (BWX5845), *smc\** SpoIIIE- (BWX5818), *smc\** MinJ- (BWX5822), and *smc\** DivIVA- (BWX5820) were spotted onto LB plates with or without 0.5 mM IPTG.

We performed a cytological characterization of *smc*,* focused on replication origin positioning and nucleoid morphology. Replication origins were visualized using TetR-CFP binding to a *tetO48* array inserted adjacent to the origin (64, 65). As can be seen in **Figure 1B**, *smc\** cells had elongated nucleoids, fewer origin foci, and occasional large origin clusters, phenotypes that indicate mild defects in both origin and bulk chromosome segregation.

### A Tn-seq screen for synthetic lethal partners of *smc\**

To identify genes that become critical when SMC function is impaired, we performed a synthetic lethal screen using Tn-seq. Using a *mariner* transposon system (66–68), deep Tn libraries were generated in both WT and *smc\** backgrounds. Genomic DNA from each library was extracted, and transposon-genome junctions were identified by deep sequencing. In the WT control, transposon insertions were found in non-essential genes, consistent with previously published results (66). In the *smc\** library, four genes were strongly depleted in Tn insertions compared with the WT library: *parB*, *spoIIIE*, *minJ*, and *divIVA* (**Fig. 1C**).

To validate these candidates, we generated isopropyl β-D-thiogalactopyranoside (IPTG)-regulated alleles of *parB*, *spoIIIE*, *minJ*, or *divIVA* (**Fig. 1D**) using three promoters of varying strengths and levels of leakiness: Phyperspank, Pspank, or Pspank* (69–71). For efficient depletion, we selected the promoter that supported cell growth that was least leaky: Pspank for *parB* and Pspank* for *spoIIIE*, *minJ*, and *divIVA*. When combined with *smc\**, all depletion strains grew in the presence of IPTG but did not form colonies without IPTG (**Fig. 1D**). We conclude that *parB, spoIIIE, minJ,* and *divIVA* become essential for growth when origin resolution and chromosome segregation are partially impaired.

### ParB is required for origin segregation in *smc\**

The synthetic lethal interaction between *smc* mutants and *parB* has been reported previously (20, 21, 60) and validates our screen. To examine the cytological consequences of this interaction, we used fluorescence microscopy to analyze origin positioning and nucleoid segregation. WT cells contained 2-4 well-separated origins distributed within segregated nucleoids (**Fig. 2A, B**). Both *smc\** and *ΔparB* single mutants showed a modest increase in cells with a single origin focus, consistent with mild origin resolution defects (**Fig. 2A, B**). However, depletion of *parB* in the *smc\** background (*smc\** ParB-) caused a strong block to origin segregation, with a sharp increase in cells containing a single origin focus and a reduced number of origins per micrometer of cell length (origin/μm) (**Fig. 2B**). This severe defect was accompanied by frequent nucleoid bisection by the division septum and an increase in anucleate cells (**Fig. 2A**, yellow carets and white asterisks, respectively), as well as elongated, heterogenous nucleoid and cell lengths (**Fig. 2A-D**). These results are consistent with previous observations in which loss of *parB* strongly exacerbates segregation defects in cells where SMC activity is reduced (20, 21).

**Figure 2.**
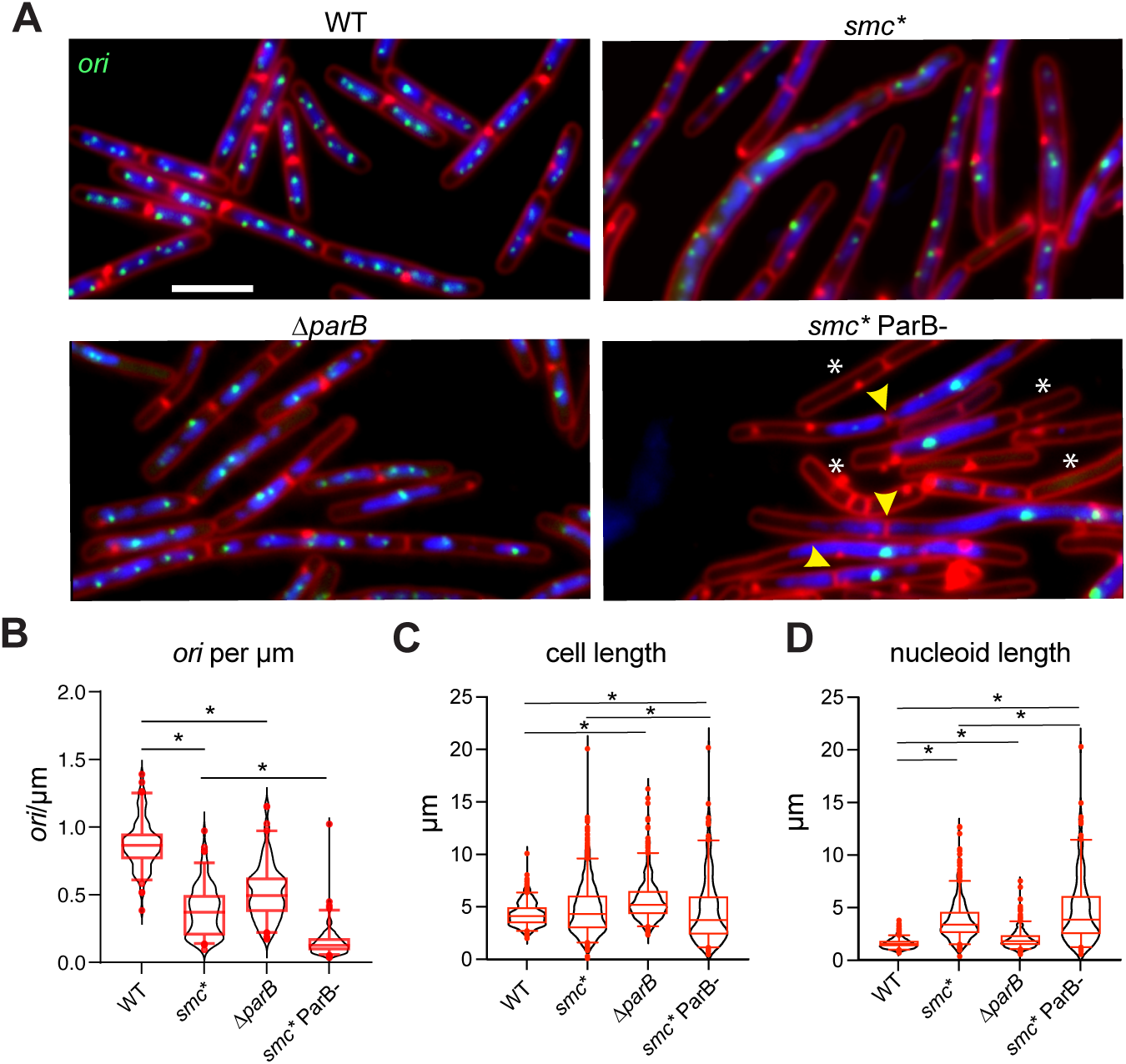
ParB is required for origin segregation in the *smc\** mutant. (A) Representative images of WT (BWX5786), *smc\** (BWX5790), *ΔparB* (BWX5839), and *smc\** ParB- (BWX5845) cells. Membranes were stained with FM4-64 (red), nucleoids were stained with DAPI (blue), and origins of replication were visualized using the *tetO*/TetR-CFP system (green). Examples of nucleoid bisection and anucleate cells are indicated by yellow carets and white asterisks, respectively. Scale bar, 4 μm. (B-D) Quantitative analyses are shown in violin plots (black). Red center lines represent the median, red boxes indicate the interquartile range, and whiskers represent 5-95 percentiles. Black asterisks denote statistical significance (*p* < 0.05) determined by the Mann-Whitney U test. (B) Origins per micrometer of cell length (*ori*/μm). 100 cells were analyzed for each strain. (C) Distribution of cell lengths. Numbers of cells analyzed: WT, 706; *smc\**, 948; *ΔparB*, 662; and *smc\** ParB-, 155. (D) Distribution of nucleoid lengths. Numbers of cells analyzed: WT, 1667; *smc\**, 561; *ΔparB*, 531; and *smc\** ParB-, 207.

### SpoIIIE prevents chromosome bisection by the division septum

Our screen identified *ΔspoIIIE* as synthetically lethal with *smc\**, consistent with previous results that *Δsmc* and *ΔspoIIIE* could not be combined by transformation (60) and with the conserved genetic interaction between SMC-family proteins and FtsK/SpoIIIE-family DNA translocases observed in other bacterial species (61–63). Unlike *smc\** ParB- cells, which exhibited severely impaired origin resolution and segregation (**Fig. 2**), *smc\** SpoIIIE- depletion cells showed only minor changes in origin positioning as measured by fluorescence microscopy and origin/μm quantification (**Fig. 3A, B**). Furthermore, we observed only modest changes in cell length and nucleoid length compared with the single mutants (**Fig. 3C, D**). By contrast, *smc\** SpoIIIE- cells exhibited a pronounced increase in chromosomes bisected by the division septum (**Fig. 3A**, yellow carets). Quantification of septum-associated chromosome bisection revealed a low percentage in WT (1.7%), *smc\** (2%), and *ΔspoIIIE* (1.6%), but ∼19% in *smc\** SpoIIIE- cells. These data are consistent with previous results (60). Since SpoIIIE/FtsK-family DNA translocases are well documented to remove DNA from the division plane and promote chromosome segregation at the terminus (31, 32, 37, 40), our results indicate that SpoIIIE becomes essential in cells impaired in chromosome segregation, facilitating terminus resolution during cytokinesis, thereby preventing chromosome bisection.

**Figure 3.**
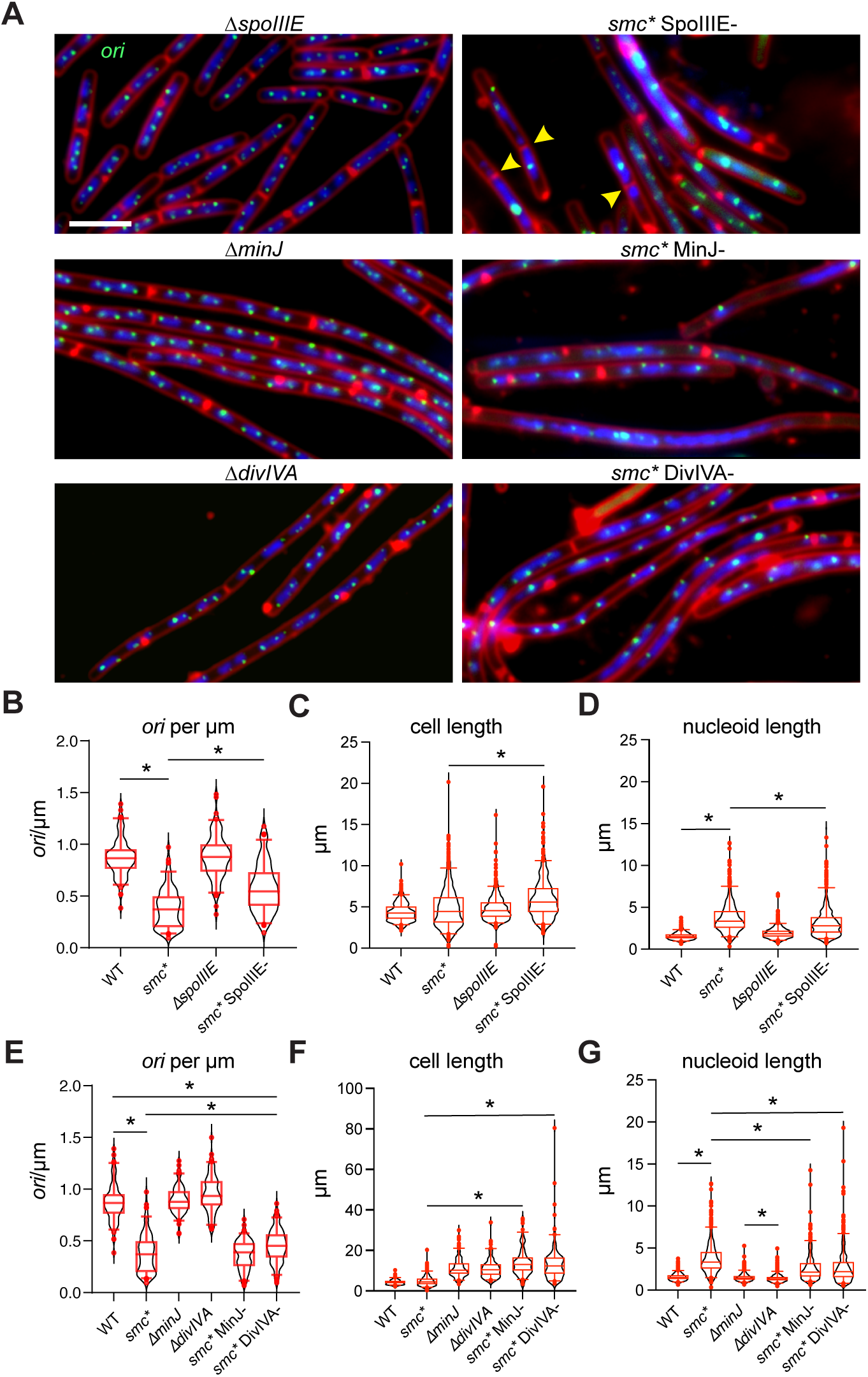
Septum positioning and chromosome clearance are essential in *smc\** cells. (A) Representative images of *ΔspoIIIE* (BWX5831), *ΔminJ* (BWX5837), *ΔdivIVA* (BWX5835), *smc\** SpoIIIE- (BWX5818), *smc\** MinJ- (BWX5822), and *smc\** DivIVA- (BWX5820) cells. Membranes were stained with FM4-64 (red), nucleoids were stained with DAPI (blue), and origins of replication were visualized using the *tetO*/TetR-CFP system (green). Examples of nucleoid bisection are indicated by yellow carets. Scale bar, 4 μm. (B-G) Quantitative analysis of representative images. WT (BWX5786) and *smc\** (BWX5790) data are the same as those shown in Figure 2. Data are shown in violin plots (black). Red center lines represent the median, red boxes indicate the interquartile range, and whiskers represent 5-95 percentiles. Black asterisks denote statistical significance (*p* < 0.05) determined by the Mann-Whitney U test. (B, E) Origins per micrometer of cell length (*ori*/μm). 100 cells were analyzed for each strain. (C, F) Distribution of cell lengths. Numbers of cells analyzed: WT, 706; *smc\**, 948; *ΔminJ*, 180; *ΔdivIVA*, 120; *smc\** MinJ-, 109; and *smc\** DivIVA-, 150. (D, G) Distribution of nucleoid lengths. Numbers of cells analyzed: WT, 1667; *smc\**, 561; *ΔminJ*, 943; *ΔdivIVA*, 728; *smc\** MinJ-, 339; and *smc\** DivIVA-, 413.

### SpoIIIE, MinJ, and DivIVA are required for terminus segregation in *smc\**

Our Tn-seq screen identified *minJ* and *divIVA* as new synthetic lethal partners of *smc\**. To characterize these interactions, we used fluorescence microscopy to analyze *smc\** MinJ*-* and *smc\** DivIVA- depletion strains. As controls, we also examined single-deletion strains (*ΔminJ* and *ΔdivIVA*), which had elongated cells as reported previously (49, 50) (**Fig. 3A, F**). In these single mutants, origin positioning and nucleoid length were modestly impaired compared with WT and were less severe than in *smc\** SpoIIIE- (**Fig. 3A, E, G**). When combined with *smc*,* both *smc\** MinJ- and *smc\** DivIVA- cells also exhibited elongated nucleoids (**Fig. 3A, F**). Although some cells displayed clustered origin foci, the defects were comparable to those observed in the *smc*\* single mutant (**Fig. 2A, E**). These observations suggest that the synthetic lethality arises from the combination of a mild origin segregation defect (*smc\**) and impaired cell division control in the absence of MinJ or DivIVA. However, these microscopy results alone were insufficient to define the underlying mechanism.

Given that *smc\** is also synthetically lethal with *ΔspoIIIE* (**Fig. 3A**) and that SpoIIIE function requires septum formation, we hypothesized that the lethality of *smc\** MinJ*-* and *smc\** DivIVA-cells resulted from the impaired SpoIIIE localization at the division septum. If correct, *smc\** SpoIIIE*-*, *smc\** MinJ-, and *smc\** DivIVA- cells should have similar chromosome segregation defects. Because origin and bulk chromosome segregation were not severely affected in *smc\** MinJ*-* and *smc\** DivIVA- cells, we next examined terminus segregation, the final step in chromosome segregation (**Fig. 4A, B**). The replication terminus was visualized with TetR-CFP binding to a *tetO48* array inserted adjacent to the terminus (65). In single mutants (*smc*, ΔspoIIIE*, *ΔminJ*, or *ΔdivIVA*), the terminus segregation pattern was similar to that of the WT, with multiple discrete foci per cell. In contrast, combining *smc\** with depletion of SpoIIIE, MinJ, or DivIVA caused severe terminus clustering, with few large foci per cell, indicative of a strong defect in the resolution and segregation of the termini (**Fig. 4A, B**). These findings argue that SpoIIIE and the MinJ/DivIVA-dependent division machinery are required to ensure proper terminus segregation in *smc\** cells.

**Figure 4.**
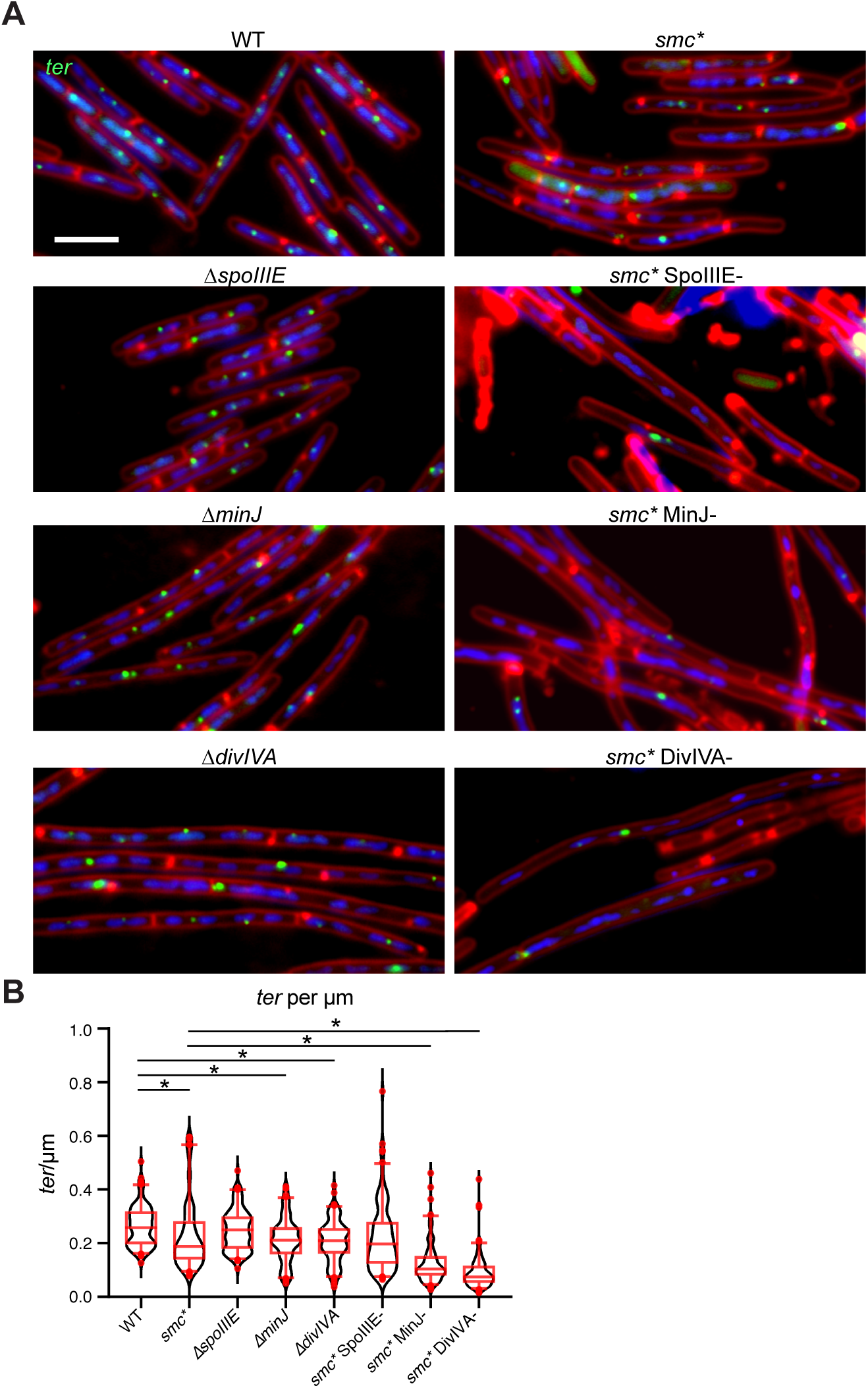
Terminus segregation requires SpoIIIE, MinJ, and DivIVA. (A) Representative images of WT (BWX6361), *ΔspoIIIE* (BWX6365), *ΔminJ* (BWX6367), *ΔdivIVA* (BWX6369), *smc\** (BWX6363), *smc\** SpoIIIE- (BWX6382), *smc\** MinJ- (BWX6377), and *smc\** DivIVA- (BWX6375) cells. Membranes were stained with FM4-64 (red), nucleoids were stained with DAPI (blue), and termini of replication were visualized using the *tetO*/TetR-CFP system (green). Scale bar, 4 μm. (B) Termini per micrometer of cell length (*ter*/μm) values are shown in violin plots (black). Red center lines represent the median, red boxes indicate the interquartile range, and whiskers represent 5-95 percentiles. Black asterisks denote statistical significance (*p* < 0.05) determined by the Mann-Whitney U test. 100 cells were analyzed for each strain.

### Synthetic lethality between MinJ/DivIVA and *smc\** is rescued by septum formation

We next asked whether the synthetic lethal interaction between *smc\** and MinJ/DivIVA arises primarily from the reduction in septum formation. Previous studies have shown that the cell division defects in *ΔminJ* or *ΔdivIVA* can be partially suppressed by deletion of *minD* (59, 72–74). Indeed, our fluorescence microscopy experiments showed that introducing *ΔminD* to *smc\** MinJ-and *smc\** DivIVA- depletion strains restored cell length, indicating the rescue of cell division defect **(Fig 5A, B)**. We then examined cell viability by spot dilution assays. In the presence of *ΔminD, smc\** MinJ- and *smc\** DivIVA- strains formed colonies regardless of IPTG addition (**Fig. 5C**). In contrast, *smc\** SpoIIIE- remained inviable upon depletion even in the *ΔminD* background (**Fig. 5C**). Because SpoIIIE activity requires an intact division septum, our results support a model in which MinJ- and DivIVA-dependent septum formation enables proper localization of SpoIIIE, which is essential for septum clearance and efficient terminus segregation to enable cell survival when SMC function is compromised.

**Figure 5.**
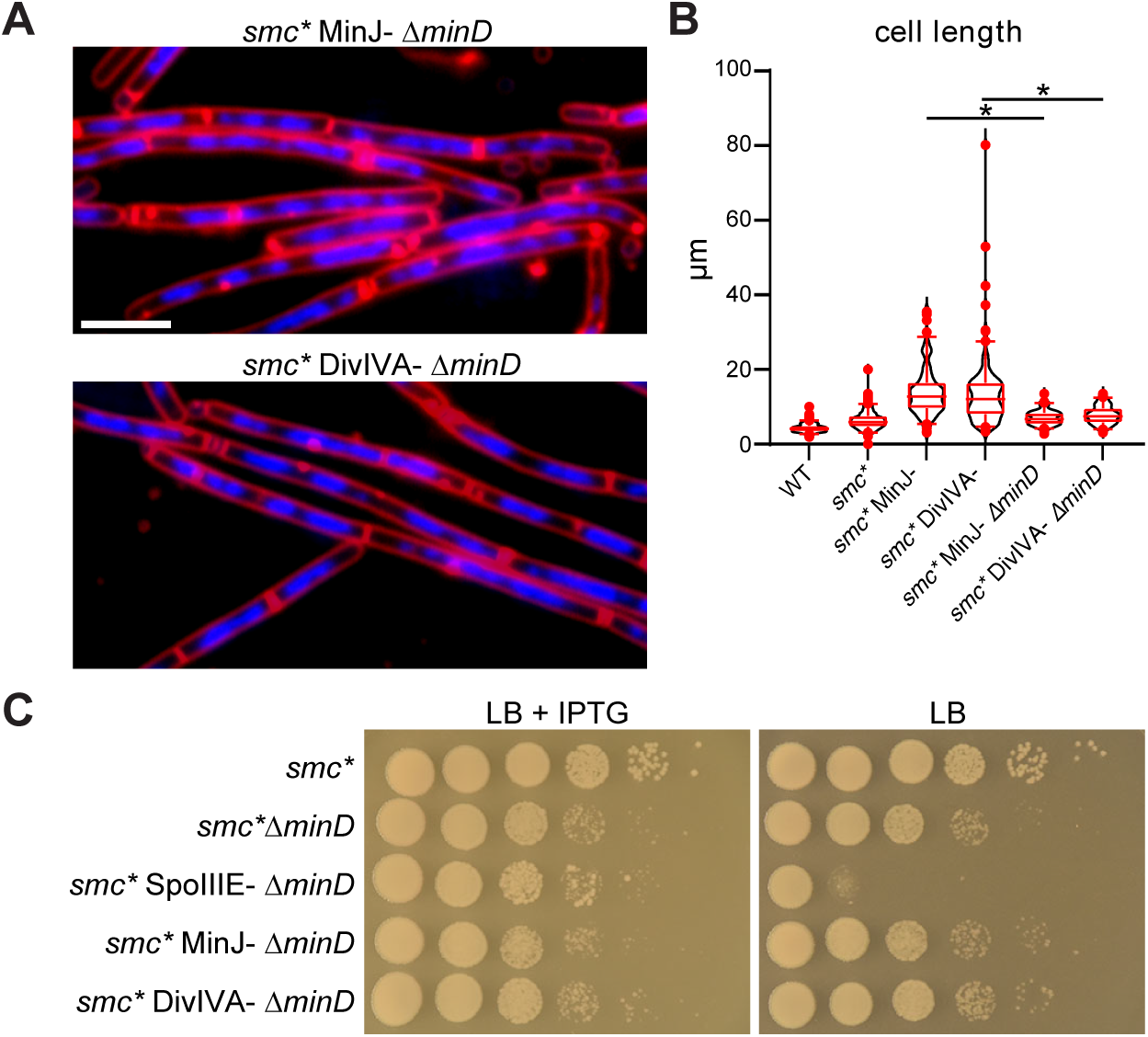
Deletion of *minD* rescues viability in *smc\** MinJ- and *smc\** DivIVA- cells. (A) Representative images of *smc\** MinJ- *ΔminD* (BWX6519) and *smc\** DivIVA- *ΔminD* (BWX6517) cells before and after depletion. Membranes were stained with FM4-64 (red), and nucleoids were stained with DAPI (blue). Scale bar, 4 μm. (B) Distribution of cell lengths is shown in violin plots (black). Red center lines represent the median, red boxes indicate the interquartile range, and whiskers represent 5-95 percentiles. Black asterisks denote statistical significance (*p* < 0.05) determined by the Mann-Whitney U test. 100 cells were analyzed for each strain. Data for *smc\** MinJ- (BW5822) and *smc\** DivIVA- (BWX5820) cells were the same as those shown in Figure 3F. (C) Serial dilutions of the indicated strains on LB agar plates with and without 0.5 mM IPTG: *smc\** (BWX2885), *smc* ΔminD* (BWX6539), *smc\** SpoIIIE- *ΔminD* (BWX6514), *smc\** MinJ- *ΔminD* (BWX6519), and *smc\** DivIVA- *ΔminD* (BWX6517).

### Septum-localized SpoIIIE promotes chromosome segregation in *smc\** cells

Previous studies showed that combining *Δsmc* with loss-of-function *spoIIIE* mutations causes synthetic sickness or lethality and that such double mutants are difficult to isolate even at 24°C (60). In addition, cells with conditional *smc* expression and *spoIIIE* mutations frequently exhibit septum-bisected chromosomes, supporting a role for SpoIIIE in translocating DNA away from the closing septum (60). Together with our findings, these observations suggest that septum-localized SpoIIIE is particularly important when SMC function is compromised. To directly examine SpoIIIE dynamics during chromosome segregation and quantify the frequency and persistence of SpoIIIE foci, we performed time-lapse fluorescence microscopy with a microfluidic device that supports chemostatic growth and long-term imaging (59) (**Fig. 6A-E**). SpoIIIE-GFP was visualized together with HBsu-mCherry to label nucleoids in WT and *smc\** cells. Images were taken every 2 min after cells were loaded into the visualization channels (**Fig. 6A**). In WT cells, SpoIIIE-GFP was distributed over the entire cell membrane, with few cells exhibiting discrete foci (**Fig. 6D**). By contrast, *smc\** cells frequently exhibited bright SpoIIIE-GFP foci positioned between segregating nucleoids, consistent with localization at the division septum (**Fig. 6E**). Quantification showed that SpoIIIE-GFP foci were detected in 33.8% of *smc\** nucleoids, compared with 10.7% in WT (**Fig. 6B**). Moreover, SpoIIIE-GFP foci persisted for an average of 60.6 min in *smc\** cells, compared with 15.7 min in WT (**Fig. 6C**). Overall, our results indicate that when SMC function is impaired, SpoIIIE is recruited to the division septum, where it translocates the trapped terminus DNA and enables completion of chromosome segregation (40). Our data further suggest that MinJ and DivIVA support this process by enabling proper septum formation, thereby providing the spatial context required for SpoIIIE-mediated DNA translocation.

**Figure 6.**
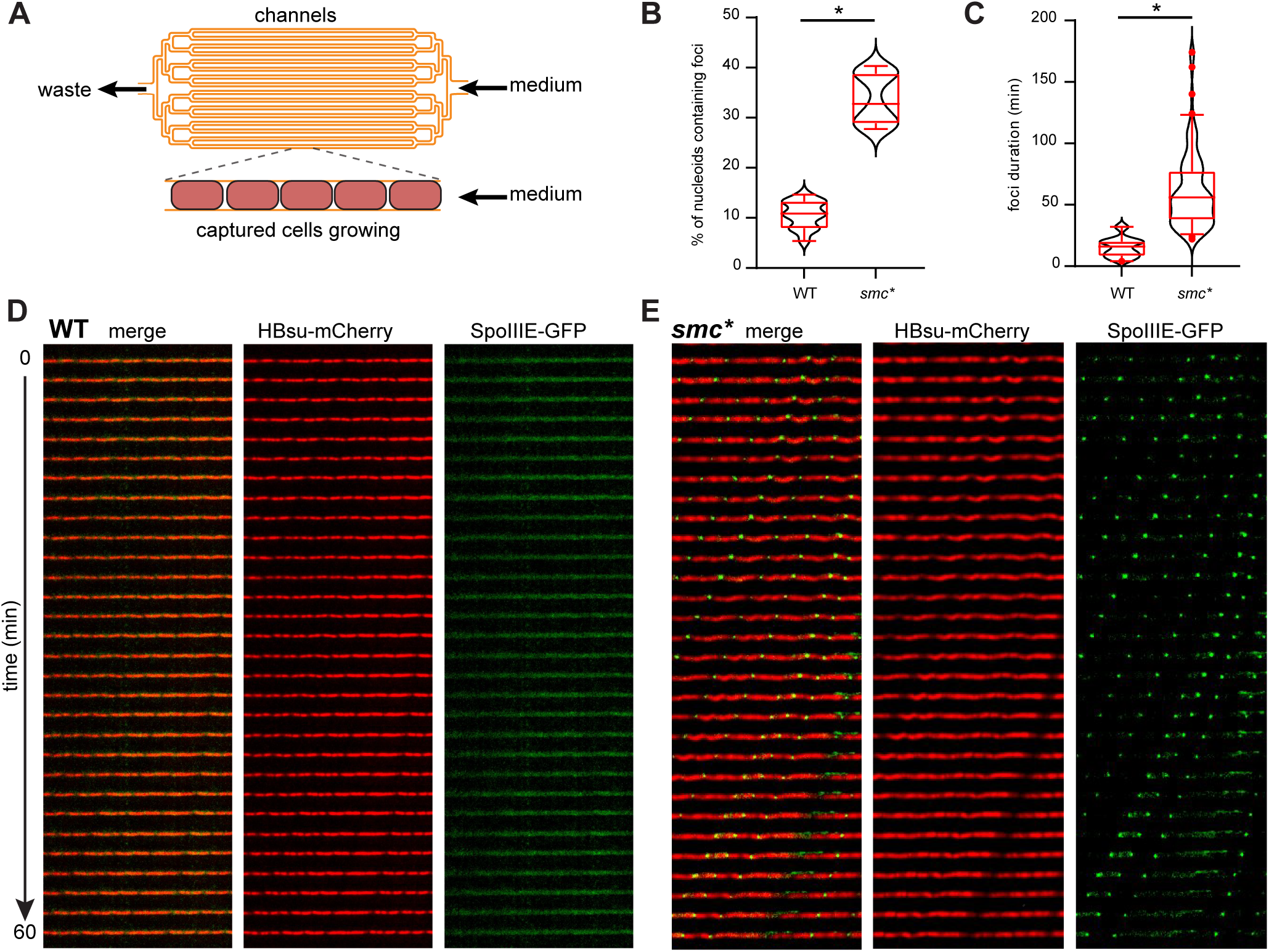
SpoIIIE-GFP forms foci at the closing septum in *smc\** cells. (A) Schematic of the microfluidic device. Cells trapped in the microchannels are grown in a chemostatic environment under a constant flow of fresh medium. (B) Quantification of the percentage of nucleoids containing SpoIIIE-GFP foci in WT (BWX5902) and *smc\** (BWX5904), for which 82,400 and 12,100 nucleoids were analyzed, respectively. Distribution is shown in violin plots (black). Red center lines represent the median, red boxes indicate the interquartile range, and whiskers represent 5-95 percentiles. Black asterisks denote statistical significance (*p* < 0.05) determined by the Mann-Whitney U test. (C) Quantification of the duration of SpoIIIE-GFP foci in WT and *smc\**, for which 20 and 100 foci were analyzed, respectively. (D-E) Kymograph analysis of WT and *smc\** cells expressing SpoIIIE-GFP (green) and HBsu-mCherry to label nucleoids (red). Overlaid images and individual fluorescence channels are shown. In each panel, cells confined within a single microfluidic channel were imaged every 2 min for 60 min to generate the kymographs. Scale bar, 4 µm.

## Discussion

In most bacteria, chromosome segregation relies on the coordinated activities of the ParABS partitioning system and SMC complexes. Previous studies showed that *smc* mutants are dependent on ParB and SpoIIIE for survival (20, 60). Here, using a systematic genetic screen, we confirmed these interactions and identified the cell division regulators MinJ and DivIVA as additional factors required when SMC function is impaired. Our results indicate that cells with reduced SMC activity become highly dependent on factors that promote origin segregation and terminus resolution.

Consistent with previous findings, ParB becomes essential for origin segregation and cell viability when SMC function is compromised (20, 60). ParB performs two separable functions: it promotes origin partitioning through ParA and loads SMC complexes at origin-proximal *parS* sites, the latter of which drives chromosome arm alignment and bulk chromosome organization (3, 8, 9, 14, 17, 18, 24, 75–77). The observation that *ΔparA* is not synthetically lethal with *smc*\* (**Fig. 1C**, top left, pink box) argues that the synthetic lethal interaction between *smc\** and *parB* is attributable to loss of ParB-mediated origin-specific loading of SMC complexes rather than ParA-dependent origin partitioning. Previous work showed that origin-specific SMC loading and chromosome arm alignment are dispensable in otherwise WT cells (14, 75). Our results indicate that these activities become important when SMC function is limited.

The requirement for SpoIIIE in *smc\** cells is also consistent with previous findings in *B. subtilis* (20, 60) and in other bacteria (61–63). Microfluidic imaging showed that SpoIIIE accumulates at midcell in *smc\** cells, consistent with recruitment to division sites containing trapped chromosomes (**Fig. 7A**). These findings indicate that defects in bulk chromosome segregation increase reliance on SpoIIIE-dependent terminus resolution (**Fig. 7A, B**).

**Figure 7.**
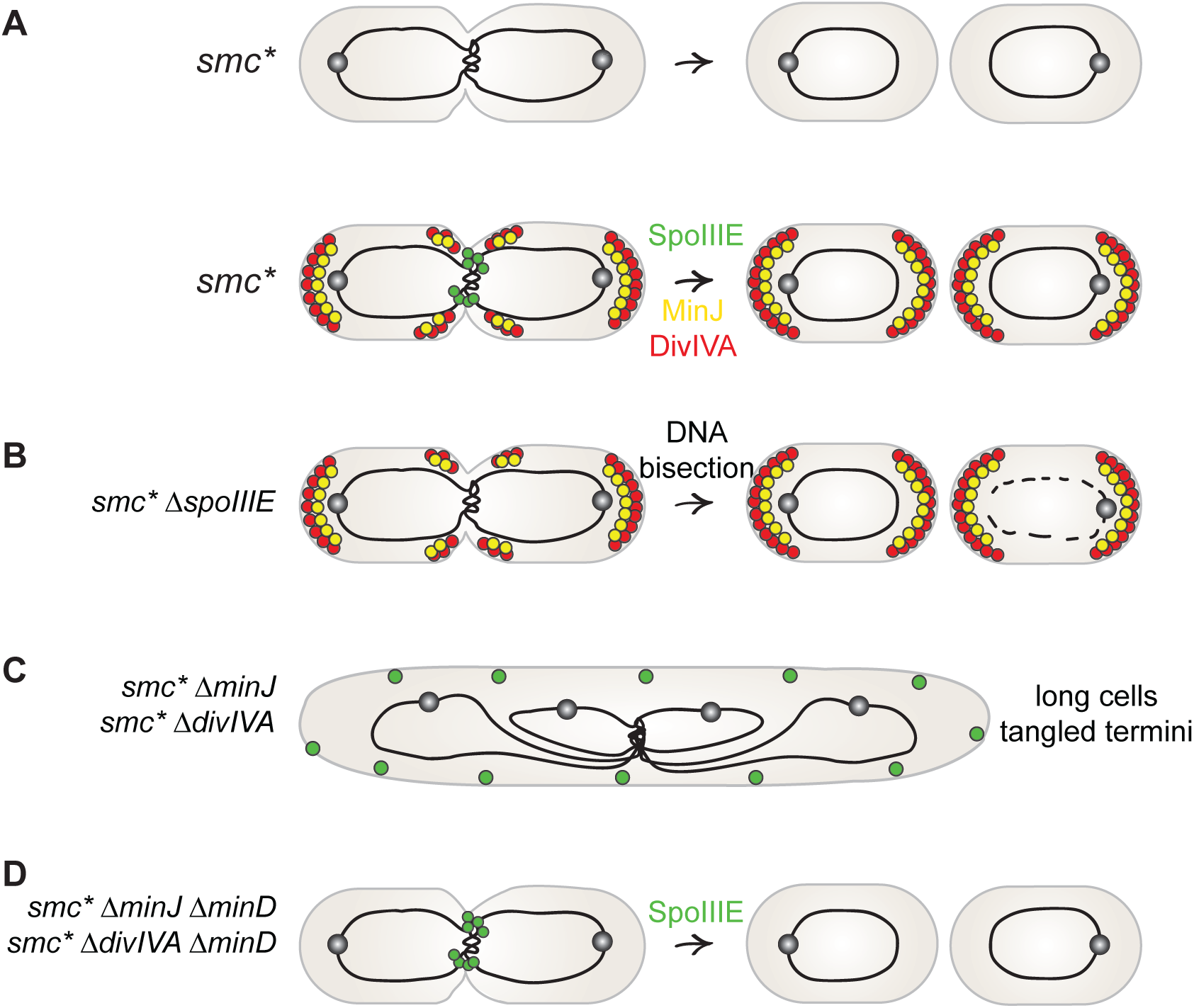
SpoIIIE and cell division are required for viability in the absence of full SMC activity. (A) The partially loss-of-function *smc\** allele impairs resolution of chromosome termini after replication, making SpoIIIE essential for completing chromosome segregation by translocating DNA away from the closing septum. Septum formation depends on MinJ and DivIVA, which provide spatial cues for SpoIIIE recruitment to resolve chromosome termini. (B) In the absence of SpoIIIE, unresolved chromosome termini remain trapped at the midcell and are ultimately bisected during septum closure, leading to DNA shearing and degradation. (C) Loss of MinJ or DivIVA in the *smc\** background disrupts septum formation, causing failure in SpoIIIE recruitment and termini resolution. (D) In the absence of MinD, cell division is restored even without MinJ or DivIVA, enabling SpoIIIE localization and rescuing cell viability.

The most notable finding from our screen was the identification of MinJ and DivIVA as factors required in *smc\** cells. Loss of either factor caused severe terminus segregation defects (**Fig. 7C**) with little effect on origin segregation, a phenotype also observed in *ΔspoIIIE*. Furthermore, deletion of *minD* restored viability, indicating that the primary role of MinJ and DivIVA in this context is to promote septum formation and thereby enable SpoIIIE function (**Fig. 7D**). These results support previous links between cell division and chromosome segregation and highlight that efficient septum formation contributes to chromosome terminus resolution when SMC-mediated segregation is compromised.

In conclusion, our results further define the pathways that become important when SMC function is reduced. We identify MinJ and DivIVA as factors that promote SpoIIIE-dependent chromosome resolution, underscoring the coordination between chromosome organization, segregation, and cell division required for faithful chromosome inheritance and cell viability.

## Materials and Methods

### General methods

All *B. subtilis* strains were derived from the *B. subtilis* 168 background (78) except as otherwise stated. Cells were grown in LB medium at 37°C. Strains, plasmids, and oligonucleotides used in this study are listed in Tables S1–S3. Strain and plasmid constructions are described in Supplemental Materials and Methods. Antibiotics were used at the following concentrations: phleomycin (Dot Scientific, DSP20200-0.025) 0.8 µg/mL, chloramphenicol (VWR, 0230-100G) 5 µg/mL, kanamycin (VWR, 0408-100G) 10 µg/mL, spectinomycin (Dot Scientific, DSS23000-25) 100 µg/mL, and erythromycin (Dot Scientific, DSE57000-50) 1 µg/mL plus lincomycin (Fisher Scientific, J61251) 25 µg/mL (79).

### Spot dilutions

For strains not requiring depletion, cells were grown in LB medium for 4 h. For depletion strains, cells were grown in LB medium supplemented with 0.5 mM IPTG for 4 h, washed three times with warm LB medium, and resuspended in LB medium. All cultures were normalized to an OD_600_ of 2, serially diluted 10-fold, and spotted onto LB plates with or without 0.5 mM IPTG. Plates were incubated at 37°C for 14 h before imaging.

### Fluorescence microscopy

Cells were grown overnight in LB medium at 22°C and then diluted into fresh LB medium and grown at 37°C to an OD_600_ of 0.2- 0.3 before imaging. Fluorescence microscopy was performed using a Nikon Eclipse Ti2 microscope with a Plan Apo 100x/1.45-numeric-aperture phase contrast oil objective. Images were captured with a scientific complementary metal-oxide semiconductor (sCMOS) camera. Membranes were stained with FM4-64 (*N*-(3-Triethylammoniumpropyl)-4-(6-(4-(Diethylamino) Phenyl) Hexatrienyl) Pyridinium Dibromide, Molecular Probes) at 3 µg/mL. Nucleoids were stained with DAPI (4’,6-diamidino-2-phenylindole, Molecular Probes) at 2 µg/mL. Images were captured and analyzed using Nikon Elements software. Final figures were prepared using Adobe Illustrator. Data analysis was performed using GraphPad Prism 10.

### Construction and isolation of *smc* hypomorphic alleles

The *smc* mutant library used in this study was generated previously (20). Briefly, the 3’ region (1.2 kb) of the *smc* gene was mutagenized by error-prone PCR using the GeneMorph II Random Mutagenesis Kit (Stratagene) with oligonucleotides oWX848 and oWX849 under conditions optimized to introduce, on average, one mutation per kilobase. Mutagenized fragments were assembled by isothermal assembly (ITA) together with three additional PCR fragments: 1) the 2.4-kb 5’ region of *smc* (amplified from the wild-type PY79 genomic DNA (80) using oWX821 and oWX847, 2) a spectinomycin-resistance cassette (amplified from pWX466 (19) using oWX438 and oWX823), and 3) a 2.2-kb downstream region of *smc* (amplified from the wild-type PY79 genomic DNA using oWX850 and oWX851). The ITA product was transformed into *B. subtilis* PY79 cells and plated on LB agar plates supplemented with 100 µg/mL spectinomycin at 30°C. Transformants were screened for reduced colony size at 37°C, and the candidate hypomorphic *smc* alleles were identified by Sanger sequencing. The chosen *smc\** contains a single point mutation V1119L and was backcrossed into the fresh PY79 WT background to generate BWX2798. For transposon sequencing, the *smc\** allele was rebuilt in the *B. subtilis* 168 background (78) with a kanamycin resistance gene, resulting in strain BWX2885. Details of strain construction can be found in the Supplemental Materials and Methods.

### Transposon mutagenesis (Tn-seq)

Tn-seq was performed following a procedure described previously (68, 79). The transposon-carrying plasmid pWX642 (68) was transformed into *smc\** (BWX2885) and WT *B. subtilis* 168. pWX642 carries a temperature-sensitive origin for *B. subtilis*, an allele of mariner Himar1 transposase, an erythromycin-resistance gene for plasmid selection, and a spectinomycin resistance gene flanked by inverted repeats detectable for the transposase. One of the inverted repeats was constructed to contain an MmeI recognition site. Transformants were selected on erythromycin plates at 30°C, then inoculated into liquid medium containing spectinomycin and grown at 22°C overnight. Cells were then plated on spectinomycin plates at 42°C to lose the plasmid. About 1 million colonies were scraped for each library.

Five OD_600_ units of cells from each pool were collected for genomic DNA (gDNA) isolation using QIAGEN DNeasy blood & tissue kit (69504). Three micrograms of gDNA were then digested using MmeI (NEB R0637S) for 90 min, and incubated with quick CIP (NEB M0525L) for 60 min at 37°C. The DNA was isolated using phenol-chloroform (1:1), precipitated using ethanol, and resuspended in 15 µL ddH_2_O. The digested DNA end was ligated to an annealed adapter (81) using T4 DNA ligase and incubated at 16°C for 16 h. The adapter-ligated DNA was then amplified with primers complementary to the adapter and the transposon inverted repeat sequence using 16-cycle PCR. The PCR product was purified using the Monarch Spin DNA Gel extraction kit (NEB T1020S) and sequenced at the IU Center for Genomics and Bioinformatics using NextSeq500. Sequencing reads were mapped to the *B. subtilis* 168 genome (NCBI NC_00964.3) and analyzed following a previously described procedure (66, 81). The results were visualized using Artemis (https://www.sanger.ac.uk/tool/artemis/) and captured by screenshots. The figures were prepared using Adobe Illustrator.

### Time-lapse microscopy in microfluidic devices

Microfluidic devices were fabricated as described previously (52). Prior to cell loading, the microchannels were coated with 1% bovine serum albumin (BSA) for 1 h to provide a passivation layer and then filled with LB medium containing 0.1% BSA. Approximately 10 μL of a saturated culture of steadily growing cells was loaded into the device through the cell reservoir and pumped into the cell-trapping region. During cell loading, a vacuum was applied to the control layer to lift the membrane and open the microchannel array, allowing cells to enter the trapping region. Once the channels were sufficiently populated with cells, positive pressure was applied to trap individual cells within the microchannels. LB medium was then pumped through the device to flush away excess cells, after which medium was supplied by gravity flow for the remainder of the experiment to maintain steady-state growth.

Cells in steady-state growth were monitored for 5 h by time-lapse microscopy. Fluorescence microscopy was performed on an Olympus IX83 microscope equipped with an Olympus UApo N 100×/1.49-NA oil-immersion objective and a Hamamatsu electron multiplying charge-coupled-device (EM-CCD) digital camera controlled with MetaMorph Advanced software. Fluorescence was excited by an Olympus U-HGLGPS fluorescence light source. Emission was collected with a tetramethylrhodamine isothiocyanate (TRITC) Semrock filter for HBsu-mCherry and a green fluorescent protein (GFP) Semrock filter for SpoIIIE-GFP. Images were captured from at least eight regions across the microchannel array at 2-min intervals. The microfluidic device was maintained at 37°C with a TC-1-100s temperature controller (Bioscience Tools). Identical microscope settings were used for all experiments to allow direct comparisons.

Following cell loading onto the microfluidic device, cells exhibited an initial adaptation period. Therefore, only data collected after steady-state behavior was established were included in subsequent analyses. Time-lapse fluorescence images were analyzed with MicrobeJ (82). Nucleoid fluorescence (HBsu-mCherry) was used to identify and track individual chromosome regions throughout the experiment. Segmentation parameters were optimized for accurate nucleoid detection and manually verified at all time points to ensure consistency. SpoIIIE-GFP localization events were identified with the particle detection function in MicrobeJ and were scored only when a fluorescent focus was positioned between two distinct nucleoids and persisted for at least six consecutive frames. This threshold was used to distinguish genuine SpoIIIE-GFP recruitment events associated with chromosome segregation from transient fluorescence signals or background noise. Measurements of nucleoid properties, SpoIIIE-GFP focus formation, and their temporal dynamics were then extracted over the course of the experiment.

## Acknowledgements

We thank Alan Grossman for the *ΔparB* strain and pKL168, the Indiana University Center for Genomics and Bioinformatics for next-generation sequencing, and the IU Nanoscience Core Facility for providing instruments. Support for this work comes from the National Institutes of Health grants R35GM145299 to D.Z.R., R35GM141922 to S.C.J., R35GM131783 to D.B.K., and R01GM141242 to X.W.

