## Supplemental Information for "Efficient septum formation is essential for chromosome segregation in *Bacillus subtilis* when SMC function is impaired"

**This PDF file includes:**

Supplemental Materials and Methods
Supplemental Tables
Supplemental References

**Supplemental Materials and Methods**

**Strain construction.**

*B. subtilis* strains were constructed by transforming plasmid DNA and genomic DNA isolated from existing strains using standard transformation assays in BMK competence medium(1).

**BWX527** [*sacA::hbsu-mCherry (cat)*] was constructed by direct transformation of a three-way ligation into *B. subtilis* PY79. The ligation consisted of the pKM064 backbone digested with EcoRI and BamHI, the *hbs* gene with its native promoter amplified from PY79 genomic DNA using primers oDR198 and oDR214 and digested with EcoRI and XhoI, and the *mCherry* gene amplified from pDR201 using primers oWX327 and oWX328 and digested with XhoI and BamHI. The ligation reaction was transformed into *B. subtilis* PY79 cells and plated on LB agar plates supplemented with 5 µg/mL chloramphenicol at 37°C. pKM064 is a cloning vector that contains *sacA (cat)*. pDR201 carries an *mCherry* gene codon-optimized for *B. subtilis* (D. Z. Rudner unpublished). The resulting strain was verified by PCR amplification of genomic DNA using primers oDR198 and oWX328, followed by Sanger sequencing using the same primers.

**BWX2798** [PY79, *smc* (spec)*] was generated by backcrossing the *smc** allele isolated from the mutant library described in the section "Construction and isolation of *smc* hypomorphic alleles" into a fresh WT background. The resulting strain contains a single point mutation V1119L and a spectinomycine resistance gene.

**BWX2883** [Bs168, *trpC2*, *smc (kan)*] was constructed by direct transformation of an isothermal assembly product into Bs168 (2). The isothermal reaction contained three PCR fragments: 1) *smc* (amplified from the BS168 genomic DNA using oWX821 and oWX849); 2) *loxP kan* (amplified from the pWX470 (3) plasmid using oWX438 and oWX823); and 3) downstream of *smc* (amplified from the Bs168 genomic DNA using oWX850 and oWX851). The ITA product was transformed into *B. subtilis* 168 cells and plated on LB agar plates supplemented with 10 µg/mL kanamycin at 37°C. The *smc* gene was sequenced using oWX523, oWX848, oWX1194, oWX1195 and oWX1196 and oWX1624.

**BWX2885** [Bs168, *trpC2*, *smc* (kan)*] was constructed by direct transformation of an isothermal assembly product into Bs168 (2). The isothermal reaction contained three PCR fragments: 1) the 3.6-kb region containing *smc** (amplified from BWX2798 genomic DNA using oWX821 and oWX849), 2) a kanamycin-resistance cassette (amplified from pWX470 (3) using oWX438 and oWX823), and 3) a 2.2-kb downstream region of *smc* (amplified from the B*. subtilis 168* genomic DNA using oWX850 and oWX851). The ITA product was transformed into *B. subtilis* 168 cells and plated on LB agar plates supplemented with 10 µg/mL kanamycin at 37°C. The *smc** gene was sequenced using oWX523, oWX848, oWX1194, oWX1195 and oWX1196 and oWX1624.

**Plasmid construction.**

**pDR181** [*spoIIIE-gfp (kan)*] was built by ligating two DNA fragments: 1) pKL168 digested by EcoRI and XhoI, yielding a plasmid backbone containing *gfp* and a kanamycin-resistance cassette; 2) the C-terminus of *spoIIIE* amplified from PY79 genomic DNA using primers oDR238 and oDR239 and digested by EcoRI and XhoI. The resulting plasmid was integrated into the PY79 chromosome by single-crossover integration. The genomic DNA from the resulting strain was transformed into the *B. subtilis* 168 background by double backcrossing. pKL168 (Lemon and Grossman, unpublished) is a Campbell-type integration vector essentially identical to pKL147 (4), except that it carries kanamycin rather than spectinomycin resistance.

**pWX209** [*pelB::tetO48 (cat)*] was built by a ligation reaction containing two DNA fragments: 1) pKM020 digested by EcoRI and HindIII to give *pelB::cat*; 2) pLAU29 (5) digested by EcoRI and HindIII to give *tetO48*. pKM020 is a cloning vector containing *pelB (cat)* (6).

**pWX649** [*yvbJ::Pspank* (optRBS) spoIIIE (spec)*] was constructed by a ligation reaction containing two DNA fragments: 1) pMS051 digested by XmaI and SpeI to give *yvbJ::Pspank* (spec)*; 2) *spoIIIE* region amplified from PY79 using oWX1167 and oWX1168 and then digested by XmaI and SpeI. pMS051 is a cloning vector that contains *yvbJ::Pspank* (spec)* (D. Z. Rudner unpublished). The construct was sequenced using oWX486 and oWX487.

**pWX659** [*yvbJ::Pspank* (optRBS) divIVA (spec)*] was constructed by a ligation reaction containing two DNA fragments: 1) pMS051 digested by XmaI and SpeI to give *yvbJ::Pspank* (spec)*; 2) *divIVA* region amplified from PY79 using oWX1188 and 1189 and then digested by XmaI and SpeI. pMS051 is a cloning vector that contains *yvbJ::Pspank* (spec)* (D. Z. Rudner unpublished). The construct was sequenced using oWX486 and oWX487.

**pWX660** [*yvbJ::Pspank* (optRBS) minJ (spec)*] was constructed by a ligation reaction containing two DNA fragments: 1) pMS051 digested by XmaI and SpeI to give *yvbJ::Pspank**

*(spec)*; 2) *minJ* region amplified from PY79 using oWX1190 and oWX1191 and then digested by XmaI and SpeI. pMS051 is a cloning vector that contains *yvbJ::Pspank* (spec)* (D. Z. Rudner unpublished). The construct was sequenced using oWX486 and oWX487.

**pWX723** [*yvbJ::Pspank (optRBS) parB ∆parS erm*] was constructed by a ligation reaction containing two DNA fragments: 1) pMS044 digested by XmaI and NheI] to give *yvbJ::Pspank (erm)*; 2) the *(optRBS) parB ∆parS* region was amplified from pWX589 (7) using oWX1668 and oWX999 and then digested by XmaI and NheI. pMS044 is a cloning vector that contains *yvbJ::Pspank (erm)* (D. Z. Rudner unpublished). The construct was sequenced using oWX486 and oDR829.

**Table S1. Strains used in this study.**

| Strain | Genotype | Reference | Fig. |
|---|---|---|---|
| BWX5786 | Bs168, *trpC2*, *smc (kan), ycgO::PftsW tetR-cfp (phleo), yycR::tetO48 (cat)* | This study | 1A, 1B, 1D, 2A-D, 5A |
| BWX5790 | Bs168, *trpC2*, *smc* (kan), ycgO::PftsW tetR-cfp (phleo), yycR::tetO48 (cat)* | This study | 1A, 1B, 1D, 2A-D, 5A |
| BWX5839 | Bs168, *trpC2*, *smc (kan), ycgO::PftsW tetR-cfp (phleo), yycR::tetO48 (cat), ΔparB (spec)* | This study | 1D, 2A-D |
| BWX5831 | Bs168, *trpC2*, *smc (kan), ycgO::PftsW tetR-cfp (phleo), yycR::tetO48 (cat), ΔspoIIIE (erm)* | This study | 1D, 3A-D, |
| BWX5837 | Bs168, *trpC2*, *smc (kan), ycgO::PftsW tetR-cfp (phleo), yycR::tetO48 (cat), ΔminJ (erm)* | This study | 1D, 3A-D, |
| BWX5835 | Bs168, *trpC2*, *smc (kan), ycgO::PftsW tetR-cfp (phleo), yycR::tetO48 (cat), ΔdivIVA (erm)* | This study | 1D, 3A-D, |
| BWX5845 | Bs168, *trpC2*, *smc* (kan), ycgO::PftsW tetR-cfp (phleo), yycR::tetO48 (cat), yvbJ::Pspank (optRBS) parB ΔparS (erm), ΔparB (spec)* | This study | 1D, 2A-D |
| BWX5818 | Bs168, *trpC2*, *smc* (kan), ycgO::PftsW tetR-cfp (phleo), yycR::tetO48 (cat), yvbJ::Pspank* (optRBS) spoIIIE (spec), ΔspoIIIE (erm)* | This study | 1D, 3A-D, |
| BWX5822 | Bs168, *trpC2*, *smc* (kan), ycgO::PftsW tetR-cfp (phleo), yycR::tetO48 (cat), yvbJ::Pspank* (optRBS) minJ (spec), ΔminJ (erm)* | This study | 1D, 3A-D, 5A |
| BWX5820 | Bs168, *trpC2*, *smc* (kan), ycgO::PftsW tetR-cfp (phleo), yycR::tetO48 (cat), yvbJ::Pspank* (optRBS) divIVA (spec), ΔdivIVA (erm)* | This study | 1D, 3A-D, 5A |
| BWX6361 | Bs168, *trpC2*, *smc (kan), ycgO::PftsW tetR-cfp (phleo), pelB::tetO48 (cat)* | This study | 4A, 4B |
| BWX6365 | Bs168, *trpC2*, *smc (kan), ycgO::PftsW tetR-cfp (phleo), pelB::tetO48 (cat), ΔspoIIIE (erm)* | This study | 4A, 4B |
| BWX6367 | Bs168, *trpC2*, *smc (kan), ycgO::PftsW tetR-cfp (phleo), pelB::tetO48 (cat), ΔminJ (erm)* | This study | 4A, 4B |
| BWX6369 | Bs168, *trpC2*, *smc (kan), ycgO::PftsW tetR-cfp (phleo), pelB::tetO48 (cat), ΔdivIVA (erm)* | This study | 4A, 4B |
| BWX6363 | Bs168, *trpC2*, *smc* (kan), ycgO::PftsW tetR-cfp (phleo), pelB::tetO48 (cat)* | This study | 4A, 4B |
| BWX6382 | Bs168, *trpC2*, *smc* (kan), ycgO::PftsW tetR-cfp (phleo), pelB::tetO48 (cat), yvbJ::Pspank* (optRBS) spoIIIE (spec), ΔspoIIIE (erm)* | This study | 4A, 4B |
| BWX6377 | Bs168, *trpC2*, *smc* (kan), ycgO::PftsW tetR-cfp (phleo), pelB::tetO48 (cat), yvbJ::Pspank* (optRBS) minJ (spec), ΔminJ (erm)* | This study | 4A, 4B |
| BWX6375 | Bs168, *trpC2*, *smc* (kan), ycgO::PftsW tetR-cfp (phleo), pelB::tetO48 (cat), yvbJ::Pspank* (optRBS) minJ (spec), ΔdivIVA (erm)* | This study | 4A, 4B |
| BWX6519 | Bs168, *trpC2*, *smc* (kan), yvbJ::Pspank* (optRBS) minJ (spec), ΔminJ (erm), ΔminD (cat)* | This study | 5A-C |
| BWX6517 | Bs168, *trpC2*, *smc* (kan), yvbJ::Pspank* (optRBS) divIVA (spec), ΔdivIVA (erm), ΔminD (cat)* | This study | 5A-C |

| | | | |
|---|---|---|---|
| BWX2885 | Bs168, *trpC2*, *smc* (kan)* | This study | 5C |
| BWX6539 | Bs168, *trpC2*, *smc* (kan), ΔminD (cat)* | This study | 5C |
| BWX6514 | Bs168, *trpC2*, *smc* (kan), yvbJ::Pspank* (optRBS) spoIIIE (spec), ΔspoIIIE (erm), ΔminD (cat)* | This study | 5C |
| BWX5902 | Bs168, *trpC2*, *sacA::hbsu-mCherry b.s. (cat), spoIIIE-gfp (kan), smc loxP (spec)* | This study | 6B, 6C, 6D |
| BWX5904 | Bs168, *trpC2*, *sacA::hbsu-mCherry b.s. (cat), spoIIIE-gfp (kan), smc* (spec)* | This study | 6B, 6C, 6E |
| **Strains that are used for strain building** | | | |
| AG1468 | *ΔparB (spec)* | (8) | |
| Bs168 | *trpC2* | (2) | |
| BKO215 | Bs168, *trpC2, ΔdivIVA (erm)* | (1) | |
| BKO220 | Bs168, *trpC2, ΔspoIIIE (erm)* | (1) | |
| BKO3053 | Bs168, *trpC2, ΔminJ (erm)* | (1) | |
| BWX527 | *sacA::hbsu-mCherry (cat)* | This study | |
| BWX811 | *ycgO::PftsW tetR-cfp (phleo), yycR::tetO48 (cat)* | (9) | |
| BWX2080 | *smc (spec)* | (10) | |
| BWX2798 | *smc* (spec)* | This study | |
| BWX2883 | Bs168, *trpC2*, *smc (kan)* | This study | |
| BWX2885 | Bs168, *trpC2*, *smc* (kan)* | This study | |
| DK7279 | *ΔminD (cat)* | (11) | |

**Table S2. Plasmids used in this study.**

| Plasmid | Description | Reference |
|---|---|---|
| pDR181 | *spoIIIE-gfp (kan)* | This study |
| pDR201 | *mCherry b.s.* | D. Z. Rudner unpublished |
| pLAU29 | *tetO48* | (5) |
| pKL168 | *gfp (kan)* | Lemon and Grossman, unpublished |
| pKM020 | *pelB::cat* | (6) |
| pMS051 | *yvbJ:: Pspank* (spec)* | D. Z. Rudner unpublished |
| pMS044 | *yvbJ::Pspank (erm)* | D. Z. Rudner unpublished |
| pWX209 | *pelB::tetO48 (cat)* | This study |
| pWX466 | *loxP spec loxP* | (3) |
| pWX470 | *loxP kan loxP* | (3) |
| pWX589 | *yvbJ::Pspank-gfp-parB (ΔparS) (cat)* | (7) |
| pWX642 | *pACYC TnKRM (spec) (amp) MmeI modified* | (12) |
| pWX649 | *yvbJ::Pspank* (optRBS) spoIIIE (spec)* | This study |
| pWX659 | *yvbJ::Pspank* (optRBS) divIVA (spec)* | This study |
| pWX660 | *yvbJ::Pspank* (optRBS) minJ (spec)* | This study |
| pWX723 | *yvbJ::Pspank (optRBS) parB ΔparS (erm)* | This study |

**Table S3. Oligonucleotides used in this study.**

| Oligos | Sequence | Use |
|---|---|---|
| oDR198 | gccctcgagttttccggcaactgcgtctttaagcgc | BWX527 |
| oDR214 | gccgaattcaaatcaccttaaatccttgacgagc | BWX527 |
| oDR238 | gccgaattctggcgctgagaagcttcttggccg | pDR181 |
| oDR239 | cggctcgagagaagagagctcatcatatttctc | pDR181 |
| oDR829 | catcaaatcttacaaatgtag | sequencing |
| oWX327 | accctcgagggatcaggacagggaccaatggtcagcaagggagaggaag | BWX527 |
| oWX328 | tatggatcccccgggctattatttgtataattcgtccattccacc | BWX527 |
| oWX438 | gaccagggagcactggtcaac | universal |
| oWX439 | tccttctgctccctcgctcag | universal |
| oWX486 | gccgctctagctaagcagaaggc | sequencing |
| oWX487 | aacggtctgataagagacaccggc | sequencing |
| oWX523 | cattcaggagtcgagattatcgctcag | sequencing |
| oWX821 | taaaatccccccttatgactcaggggg | *smc* mutant library |
| oWX823 | cagtaacgaggaaagaggttaaaagggatccttctgctccctcgctcag | *smc* mutant library |
| oWX847 | cttttcagatacggcagagagctcttc | *smc* mutant library |
| oWX848 | gaagagctctctgccgtatctgaaaag | *smc* mutant library |
| oWX849 | ctgagcgagggagcagaaggatcccttttaacctctttcctcgttactg | *smc* mutant library |
| oWX850 | gttgaccagtgctccctggtctaacgaggaaagaggttaaaagatgagc | *smc* mutant library |
| oWX851 | cgtcagcctcaagcagcgcaagacgg | *smc* mutant library |
| oWX999 | tttgctagccagagtggaggcaagaacgccttaaccc | pWX723 |
| oWX1167 | aaacccgggacataaggaggaactactatggtggcaaagaaaaaacgaaaatc | pWX649 |
| oWX1168 | cgcactagtttaagaagagagctcatcatatttctc | pWX649 |
| oWX1188 | aaacccgggacataaggaggaactactatgccattaacgccaaatgatattcac | pWX659 |
| oWX1189 | cgcactagtttattccttttcctcaaatacagcgtcgac | pWX659 |
| oWX1190 | aaacccgggacataaggaggaactactatggtgtctgttcaatggggaattgaac | pWX660 |
| oWX1191 | cgcactagtttatgatcccgaagcgactgcttcgtc | pWX660 |
| oWX1194 | gggaaagtggaagagatcctgagc | sequencing |
| oWX1195 | cttcacaatgaaaatgtcgaagag | sequencing |
| oWX1196 | gcccggcattcatcatttctcggg | sequencing |
| oWX1624 | ttcgatatcataggcagtcagcgc | sequencing |
| oWX1668 | aaacccgggacataaggaggaactactatggctaaaggccttggaaaaggg | pWX723 |
